# Synthetic sphingolipid analogs and anti-cancer drugs synergistically induced human colon and pancreatic cancer cell death through inhibiting ceramide metabolism

**DOI:** 10.1101/2021.10.01.462804

**Authors:** Jing Song, Arie Dagan

**Affiliations:** Department of Biochemistry and Molecular Biology, Hadassah Medical School, The Hebrew University of Jerusalem, Jerusalem, Israel

**Keywords:** analogues, ceramide, cell toxicity, anti-cancer drugs

## Abstract

Ceramide metabolism is a potential target for anti-cancer therapy. Studies show that chemotherapeutic agents can induce apoptosis and it is mediated by ceramide. Synthesized sphingolipid analogs can induce cell death in human lymphocytes and leukemia cells. By screening a group of synthetic sphingolipid analogs, we found that low concentrations of AD2750 and AD2646 induced cell death in human cancer cells by preventing ceramide from converting to sphingomyelin, individually or in combination with commercial cancer drugs. The combination of low concentrations of Taxol and AD2750 or AD2646 significantly increased cell death on human colon cancer cells (HT29). Co-administering low concentrations of Doxorubicin with AD2750 or AD2646 elevated cellular toxicity on human pancreatic cancer cells (CRL1687). This synergistic effect is related to the elevated cellular ceramide. Combining AD2750 or AD2646 with chemotherapy drugs can be used to manipulate ceramide and sphingomyelin metabolism, potentially to affect the growth of human cancer cells and increase the effectiveness of anti-cancer drugs on killing cancer cells.

## Introduction

Sphingolipids are essential constituents of all cell membranes. The backbone of all sphingolipids is ceramide. Ceramide levels can be elevated by inhibition of the synthesis of sphingomyelin, glycolipids, or ceramide phosphate from ceramide, or by inhibition of the hydrolysis of ceramide by ceramidases. It can also be elevated by activation of the hydrolysis of sphingomyelin, or glycolipids to ceramide; or by activation of the *de-novo* synthetic pathway of ceramide (Morad, Davis et al. (2015). Incubating cells with synthetic cell-permeable short-chain ceramides or modifying sphingolipid metabolism can increase cellular ceramide level. Previously, we used synthesis fluorescent substrates to study sphingolipid metabolism *in vitro* and in intact cells (Dagan, Agmon et al. 2000). We have shown that fluorescent derivatives of fatty acids were transported into cultured human leukemic myeloid cells and their subsequent metabolic utilization (Morand, Fibach et al. 1982).

Sphingosine and ceramide can be phosphorylated and dephosphorylated by kinase and phosphatase activities, which exist in multiple compartments. Sphingosine-1-phosphate (S1P) as well as ceramide play essential roles in cell signaling (Ogretmen 2018). S1P stimulates cell survival, proliferation, and tissue regeneration. Ceramide has been shown to be involved in stress-related cellular responses and apoptosis, S1P can be degraded by S1P lyase which is located at the cytoplasmic side of the ER or can be dephosphorylated by specific phosphatases at the luminal side and form ceramide. Ceramide transfer proteins (CERT) were implicated in cancer biology, and the data showed that down-regulation of CERT results in increased sensitivity against chemotherapeutic agents (Swanton, Marani et al. 2007), suggesting that alterations of sphingolipid metabolism by CERT might confer a survival advantage to cancer cells (Hannun and Obeid 2008).

Several drugs (Haimovitz-Friedman, Kan et al. 1994) elevated ceramide in treated cells. These drugs include the anthracyclines doxorubicin and daunorubicin (Carvalho, Santos et al. 2009), the vinca alkaloids vincristine and vinblastine, antiestrogens such as tamoxifen (Vethakanraj, Sesurajan et al. 2018), the novel synthetic retinoid N-(4-hydroxy phenyl) retinamide (4-HPR) (Guilbault, De Sanctis et al. 2008, Wang, Maurer et al. 2008), and the taxane paclitaxel (Taxol) (Qiu, Zhou et al. 2006). Daunorubicin induces ceramide accumulation that subsequently killed cancer cells. Several cellular stress agents elevate ceramide, initiating a cascade of events leading to arrest of the cell cycle, apoptosis and cell death (Radin 2001, Ogretmen 2006, Hannun and Obeid 2008).

In our lab, we synthesized many non-natural sphingolipid analogues (Dagan, Wang et al. 2003, Labaied, Dagan et al. 2004, Granot, Milhas et al. 2006, Song, Dagan et al. 2017). In this study, we tested several of these analogues on human cancer cell lines including human colon (HT29) and pancreatic (CRL1687) cancer cells and analyze the results by MTT test. To understand how ceramide metabolism influences cell toxicity, we developed a fluorescent procedure to quantify the biosynthesis of cellular ceramide. Furthermore, we studied whether combining synthetic sphingolipid analogues and commercial cancer drugs can increase cell toxicity in human cancer cells, and the mechanism of the synergistic effects. Here, we show that ceramide level on human colon and pancreatic cancer cells increased following incubation with the synthetic analogues AD2750 and AD2646. AD2750 and AD2646 inhibited the biosynthesis of sphingomyelin (SPM) and glycosphingolipids and elevated cellular ceramide, then induced cell death on human cancer cells (Testi 1996, Darroch, Dagan et al. 2005, Fyrst and Saba 2010). Combining AD2750 or AD2646 with Taxol or Doxorubicin can significantly increase ceramide level in both HT29 and CRL1687 cells, affect their growth and increase the effectiveness of anti-cancer drugs on killing cancer cells.

## Materials and methods

### Materials

All solvents were of analytical grade. The chemicals were purchased from Sigma Aldrich (St Louis, USA). Bodipy propanoic and dodecanoic acids were from Invitrogen (Molecular Probes, Oregon, USA). Culture media and supplements were from Biological Industries (Kibbutz Beth Ha’Emek, Israel). Taxol, Doxorubicin, Oxaliplatin, Gemcitabine, 5-FU, Vinblastine, Vincristine, and Reservatrol were from Hadassah Hospital.

### Cell culture and cell viability

Human colon cancer cells (HT29), human melanoma cells (MDA-MB-435), human lymphoblast cells (ARH77) and human pancreatic cancer cells (CRL1687) were from ATCC and maintained in RPMI-1640 with L-Glutamine and Phenol Red, supplemented with 10% (v/v) heat-inactivated Fetal Calf Serum (FCS), 15mM HEPES, 1% (v/v) penicillin-streptomycin, and 1% L-glutamine. Cell viability test was done by MTT test according to manufacturer’s protocol. Thiazolyl blue tetra-zolium bromide is from Sigma Aldrich (St Louis, USA).

### Synthesis of the analogues

AD2646, AD2730, AD2725, AD2673, AD2750 and AD2900 were all synthesized in our laboratory (Dagan, Wang et al. 2003, Granot, Milhas et al. 2006, Song, Dagan et al. 2017). All compounds were dissolved in DMSO and kept at −80°C until they were tested. Then, the compounds were diluted by RPMI1640 (without L-Glutamine and without Phenol Red) supplemented with 10% (v/v) heat-inactivated FCS at different concentrations.

### Synthesis of fluorescent ceramides

Bodipy-C12-Ceramide(B12Cer)(Dagan, Wang et al. 2003), AD2915 (CH3(CH2)12-CH-CH-CHOH-CH-[NHCO-(CH2)11-Bodipy]-CH2OH): 1µmol bodipy dodecanoic acid was mixed with 1.2µmol of sphingosine in dimethyl formamide, 1.2µmol of 1-hydroxybenzotriazol and 2µmol 1-(3-dimethylaminopropyl)-3-ethylcarbodiimide (EDAC). The mixture was stirred overnight at room temperature, the solvent was evaporated and the product, i.e., Bodipy-C12-Ceramide, was purified on a silica gel column and product 850µmol (85% yield). The structure and purity of each of the synthetic products was analyzed and confirmed by nuclear magnetic resonance (NMR) and mass spectrometry (Dagan, Wang et al. 2003, Granot, Milhas et al. 2006, Song, Dagan et al. 2017).

### *In situ* assays

#### Inhibition of Bodipy-C12-ceramide (B12Cer) hydrolysis

HT29 cells were incubated for 24 h with fresh medium at 37°C. Then, the synthetic compounds or commercial drugs were added at various concentrations. After 30min, B12Cer (final concentration of 2-3nmol/ml) was added. The cells were incubated for additional 24-72h and then harvested by trypsinization, centrifuged at 2,000g, and the pellets were resuspended in phosphate-buffered saline (PBS). Aliquots of cells were discarded, and the remaining sample was processed as follows. Cells were centrifuged at 2,000g, and the cell pellet was resuspended in 1ml of chloroform-methanol (1:2, v/v). The solvent was collected, and the cell pellet was resuspended in 1ml of chloroform-methanol (1:1, v/v). The resulting solvent was evaporated, and the dried lipid extract was dissolved in 200µl of ethanol. The fluorescent products, B12Cer, B12SPM, and B12GC were then detected and quantified by HPLC.

#### High-performance liquid chromatography (HPLC) system

The HPLC system consisted of Waters 2690 delivery system, Waters Millennium software, Waters 474 scanning Fluorescence detector, using RP select B 125-4 Column (Merck). The chromatography was carried out at a flow rate of 1.5ml/min starting with a mobile phase of 80% methanol and 20% H2O for 3 minutes, then the composition was changed to 90% methanol and 10% H2O at the 4th minute and this ratio was kept during the next 4 minutes. At the 8^th^ minute, the composition was gradually changed to 60% methanol and 40% Dichloromethane for 3 minutes. The fluorescent derivatives were monitored at an excitation wavelength of 505nm and an emission wavelength of 530nm. Quantification of all the sphingolipid peaks was calculated by Waters Millennium software.

#### Statistical analysis

Results are expressed as Mean± SEM of at least 3 values per experiment. Each experiment was repeated at least 3 times. Mean values were compared using the Kruskal-Wallis test or the Holm-Sidak method, with alpha =0.05 using Prism software. Differences were considered statistically significant when P <0.05.

#### Consent to Participate (Ethics)

No human or animal samples were used in this study.

## Results

### Synthetic sphingolipid analogues and commercial anti-cancer drugs induced cytotoxicity on human cancer cells

To determine the proper concentrations and incubation time of synthetic sphingolipid analogues and anti-cancer drugs on human cancer cells, the half maximal inhibitory concentrations (IC50) of HT29 cells treated with 6 synthetic sphingolipid analogues (Figure1 and table1 A) and commercial drugs, including Taxol, Doxorubicin, 5-FU, Vinblastine, Vincristine, Reservatrol (table1B) were accessed by MTT test. IC50 of all analogues were lower on longer incubation times. This indicates analogues may be effective at a longer incubation time than 24h. The IC50 of Taxol on HT29 cells is under 10μM after 72h incubation. The concentrations which represent 10% cytotoxicity (IC10) to HT29 cells for different compounds were selected as the initial concentrations for the combinatorial experiments. All analogues were also tested on ARH77, MDA-MB-435 and CRL1687 cells (data not shown).

**Figure 1.**
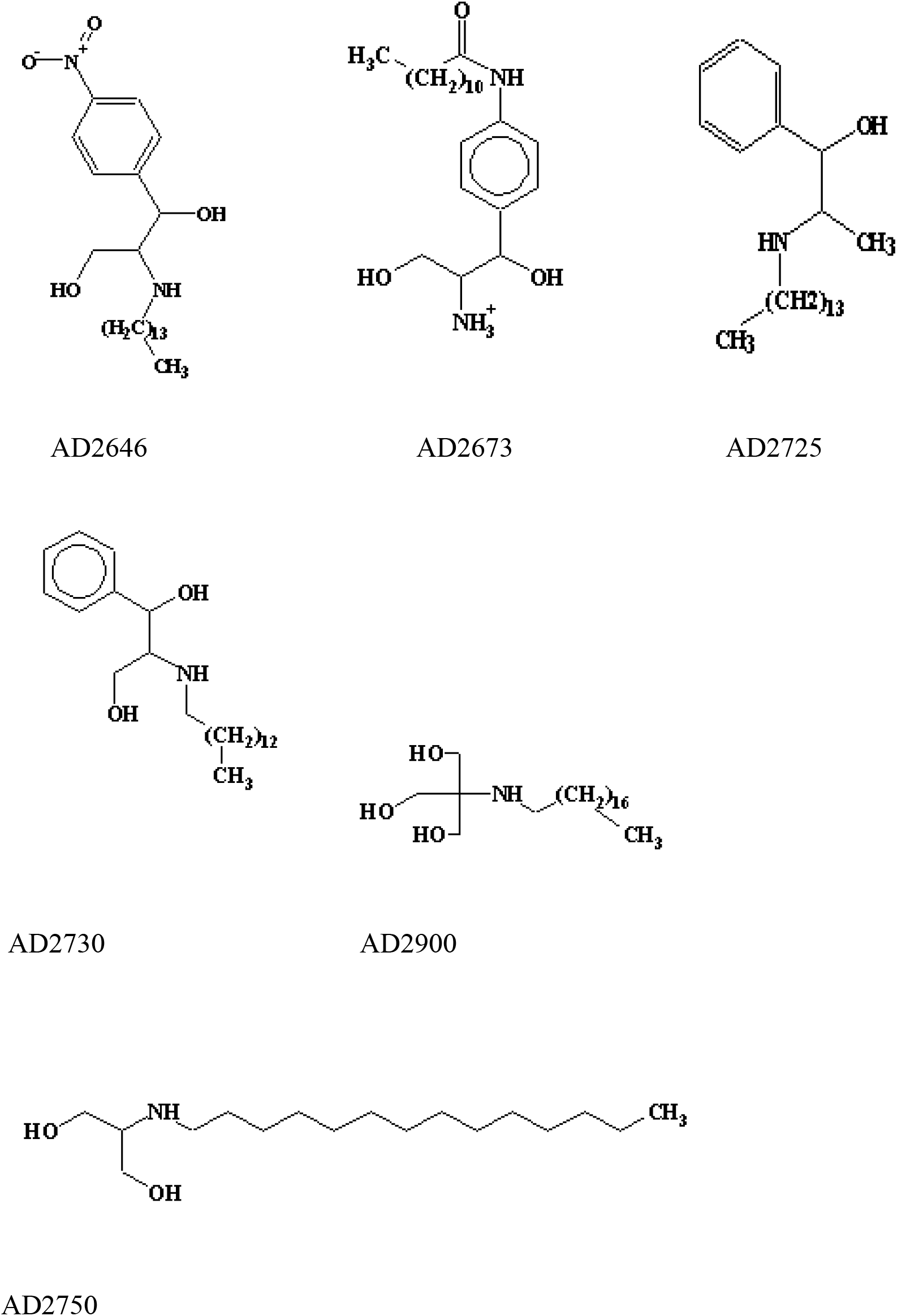
Structure of synthetic sphingolipid analogues.

**Table 1.** IC50 (μM) of (A) synthetic sphingolipid analogues and (B) commercial drugs on human colon cancer cells (HT29) in MTT test. Experiments were done for 24, 48 or 72 h. Each experiment was repeated at least 3 times. NA stands for not done.

| IC50(μM) | 24h | 72h |
| --- | --- | --- |
| AD2750 | 7 | 5 |
| AD2900 | 20 | 5 |
| AD2646 | 10 | 5 |
| AD2730 | 7 | 6 |
| AD2725 | 8 | 7 |
| AD2673 | 25 | 15 |

**Table 1.** IC50 ( μM) of (A) synthetic sphingolipid analogues and (B) commercial drugs on human colon cancer cells (HT29) in MTT test. Experiments were done for 24, 48 or 72 h. Each experiment was repeated at least 3 times. NA stands for not done.
| IC50(μM) | 24h | 48h | 72h |
| --- | --- | --- | --- |
| Taxol | NA | 10 | NA |
| Doxorubicin | >50 | >50 | >50 |
| 5-FU | >50 | >50 | >50 |
| Reservatrol | >100 | >100 | >100 |
| Vincristine | 35 | NA | 18 |
| Vinblastine | 35 | NA | 20 |

### Synthetic sphingolipid analogues inhibited ceramide metabolism in human cancer cells

To quantify the synthesis of sphingomyelin and glycosyl-ceramide in intact cells, we synthesized a long-chain fluorescent ceramide, bodipy dodecanoyl ceramide (B12Cer). Following the incubation of B12Cer with various human cancer cells, products of its utilization for sphingolipid biosynthesis, i.e., bodipy-C12-sphingomyelin (B12SPM), bodipy-C12-β-glucosylsphingosine and/or β-galactosylsphingosine (B12GC) were quantified in cell lysates by HPLC. Retention times of Bodipy-C12-conjugated fatty acid (B12FA), B12Cer and B12SPM standards are at 2.5, 5.9, 6.5 and 8.2min, respectively. We also examined the difference on the synthesis of B12SPM and B12GC after 72h of incubation of various human cancer cell lines labeled with B12Cer. B12SPM accumulated more than 3-fold on HT29 than on other human cancer cells (Fig 2). This suggests that HT29 cells could be used as a model to show relative faster metabolism of sphingolipid *in vitro*, since ceramide converted more to sphingomyelin in HT29 cells compared with ARH77, MDA-MB-435 and CRL1687.

**Figure 2.**
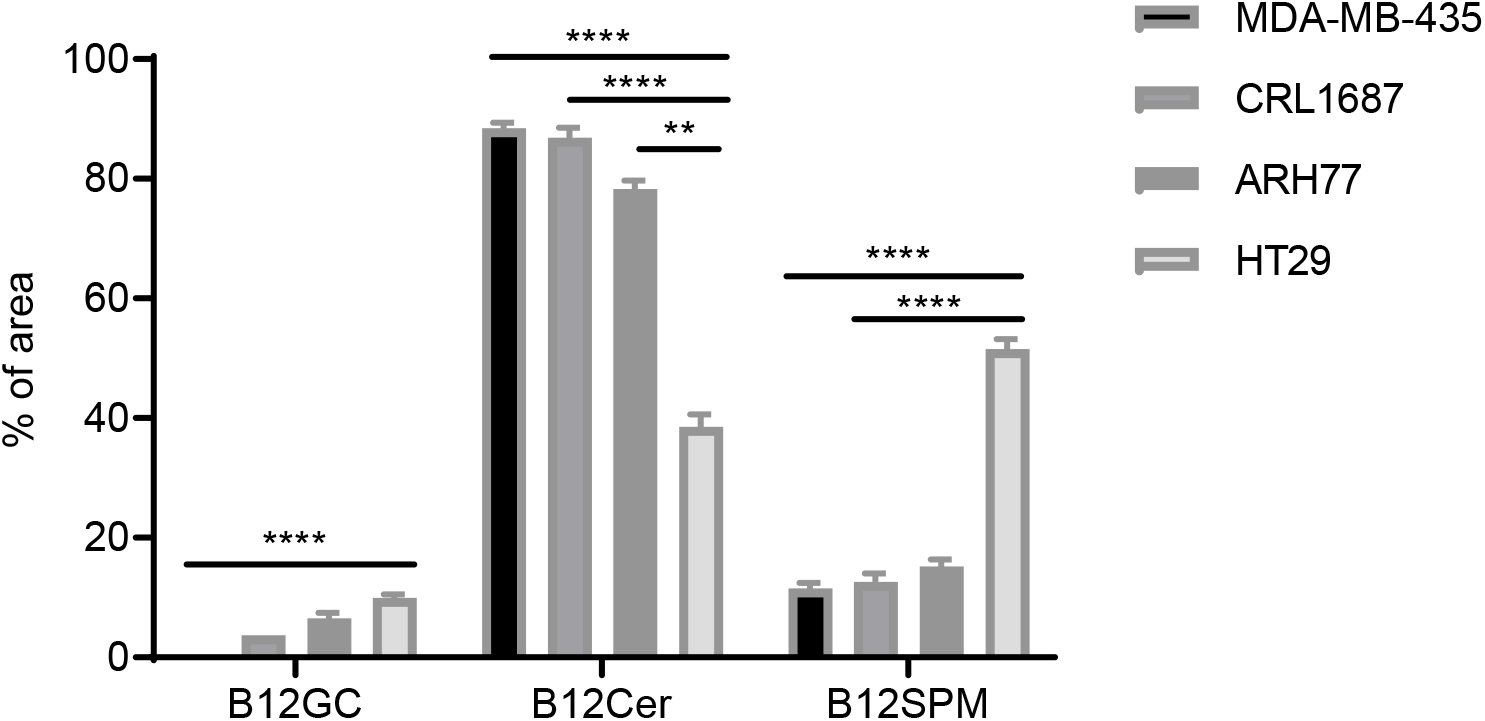
B12SPM accumulated more in HT29. Different GC/SPM contents in human cancer cell extracts including HT29, MDA-MB-435, ARH77 and CRL1687. % of total area were shown. % of area stands for the percentage of area of each curve in total area of all curves in HPLC. The total of % of area is 1. Statistical significance determined using the Holm-Sidak method, with alpha = 0.05. Each experiment was repeated at least 3 times (Mean± SEM). (*P<0.05, **P<0.01, ****P<0.0001.)

To select the right timepoint to work with, we tested the conversion of B12Cer in HT29 cells lysate at different timepoints (Fig 3). The major change of ceramide metabolism happened at either 48h or 72h. The conversion of B12Cer to B12GC and B12SPM significantly increased from 48h to 72h incubation. This could be due to longer incubation such as 72h may lead to many secondary responses in ceramide metabolism. In the following experiments, we tested both 48h and 72h incubation. We next investigated how the dose of sphingolipid analogues influenced the conversion of B12Cer on HT29 *in situ*. The conversion of B12Cer to B12SPM and B12GC was affected by compounds AD2750 and AD2900 in a dose-dependent manner (Fig 3B). 4µM AD2646 and 2.5µM AD2730 significantly elevated ceramide level in cancer cells. Based on these results, we have chosen the working concentration to co-administrate synthetic analogues and commercial drugs to cancer cells. Ceramide is a pro-apoptotic agent which plays a critical role in cell death. These results suggested that the increased level of ceramide could contribute to the cytotoxicity of HT29.

**Figure 3.**
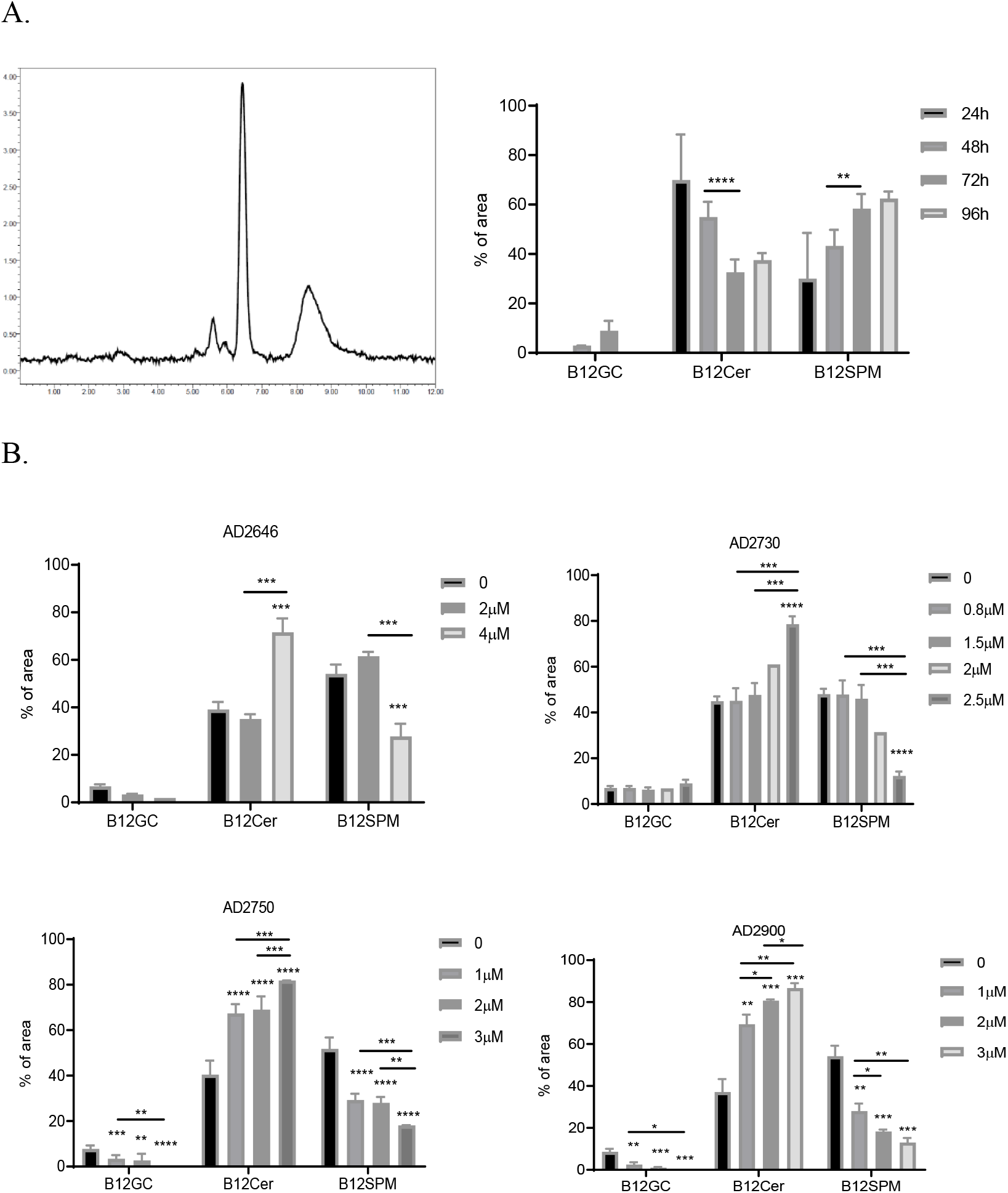
Synthesized analogues inhibited the ceramide conversion on HT29 cells in situ. A). Retention times of B12GC, B12Cer, B12SPM from HT29 cell extracts are at 5.6, 5.9, 6.5 and 8.2minute analyzed by HPLC, respectively. HT29 cells were chase-labelled by BC12Cer and incubated for 24, 48, 72 or 96 h. B). Effects on the conversion of B12Cer to B12 GC and B12SPM in HT29 extracts treated by AD2646 (2μM, 4μM), AD2730 (0.8μM, 1.5μM, 2μM, 2.5μM), AD2750 (1μM, 2μM, 3μM), AD29000 (1μM, 2μM, 3μM) are shown. HT29 cells were labelled before adding analogues to cells culture for 3 days, % of total area was shown. Statistical significance determined using the Holm-Sidak method, with alpha = 0.05. Each experiment was repeated at least 3 times (Mean± SEM). Significance was compared to 0 unless being indicated. (*P<0.05, **P<0.01, ***P<0.001, ****P<0.0001)

Then, we investigated whether combining 2 synthetic analogues can enhance cytotoxicity on cancer cells. With the co-administration of 2.5μM AD2730 and AD2900 on cancer cells, the cell viability of MDA-MB-435 (Fig. 4A) decreased significantly compared with AD2730 or AD2900 alone. Similar results were observed with HT29 cells when combining AD270 and AD2750 (Fig. 4B).

**Figure.4.**
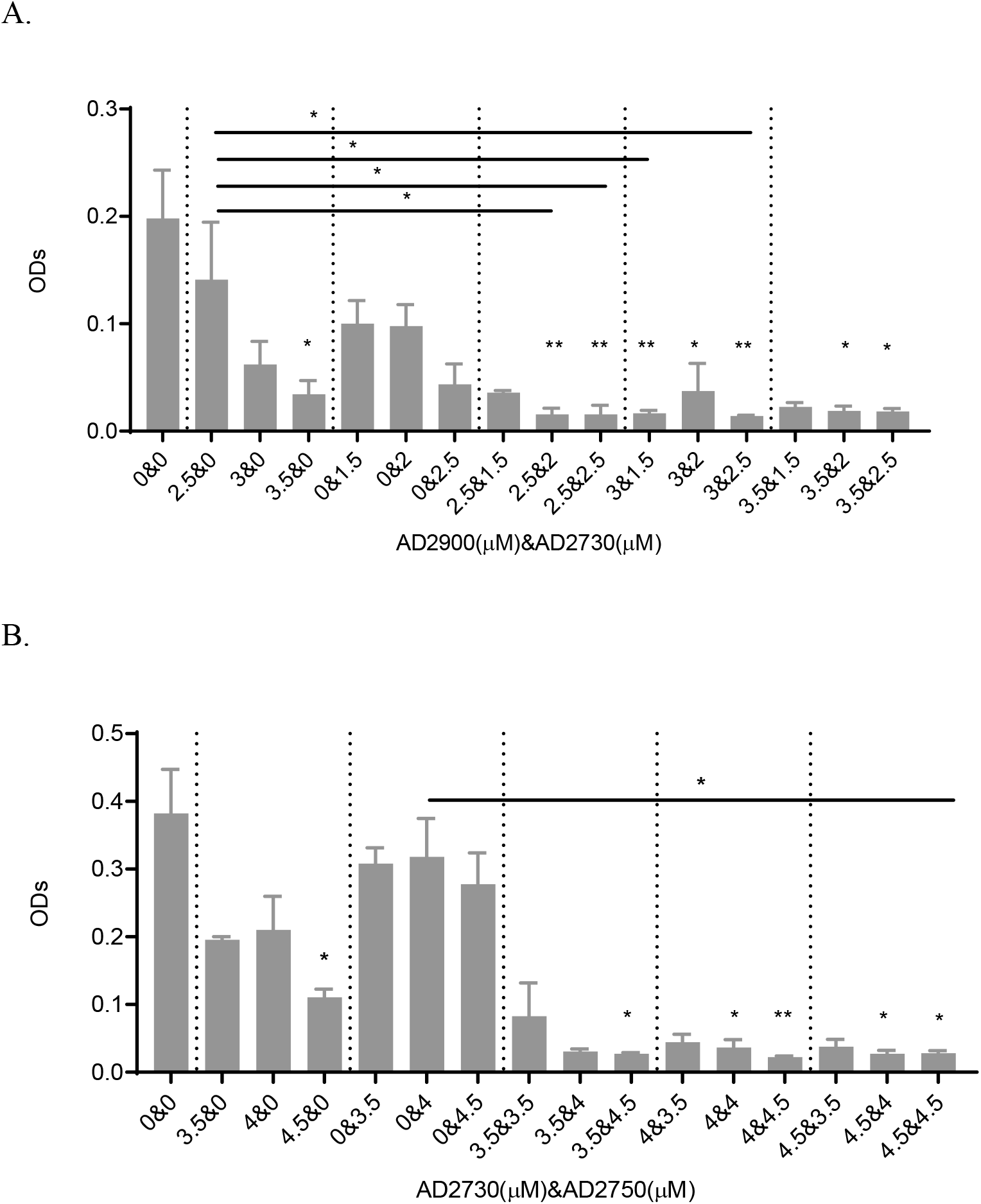
Two synthetic analogues synergistically increase cell death on human cancer cells. A). MDA-MB-435 cells were treated by the combination of AD2900 (2.5μM, 3μM, 3.5μM) and AD2730 (1.5μM, 2μM, 2.5μM) for 24 h. B). HT29 cells were treated by the combination of AD2730 (3.5μM, 4μM, 4.5μM) and AD2750 (3.5μM, 4μM, 4.5μM) for 48 h. Cell viability was checked by MTT tests. ODs were shown. Each experiment was repeated at least 3 times (Mean± SEM). Kruskal-Wallis test was used, significance was compared to 0&0 unless being indicated. (*P<0.05, **P<0.01.)

### Synergistic effects by combining Doxorubicin or Taxol with sphingolipid analogues on inducing cytotoxicity on cancer cells

Next, we investigated whether combining cancer drugs and synthetic analogues can enhance cytotoxicity on cancer cells more efficiently. When doxorubicin or AD2750 was applied to CRL1687 cells individually, cell viabilities were 71%, 62.5% and 54%, respectively. When adding both doxorubicin & AD2750 to CRK1687 cells, the cell viability dramatically decreased to 8% (Fig. 5A). Similar results were shown with the combination of Doxorubicin and AD2646 (Fig. 5B).

**Figure.5.**
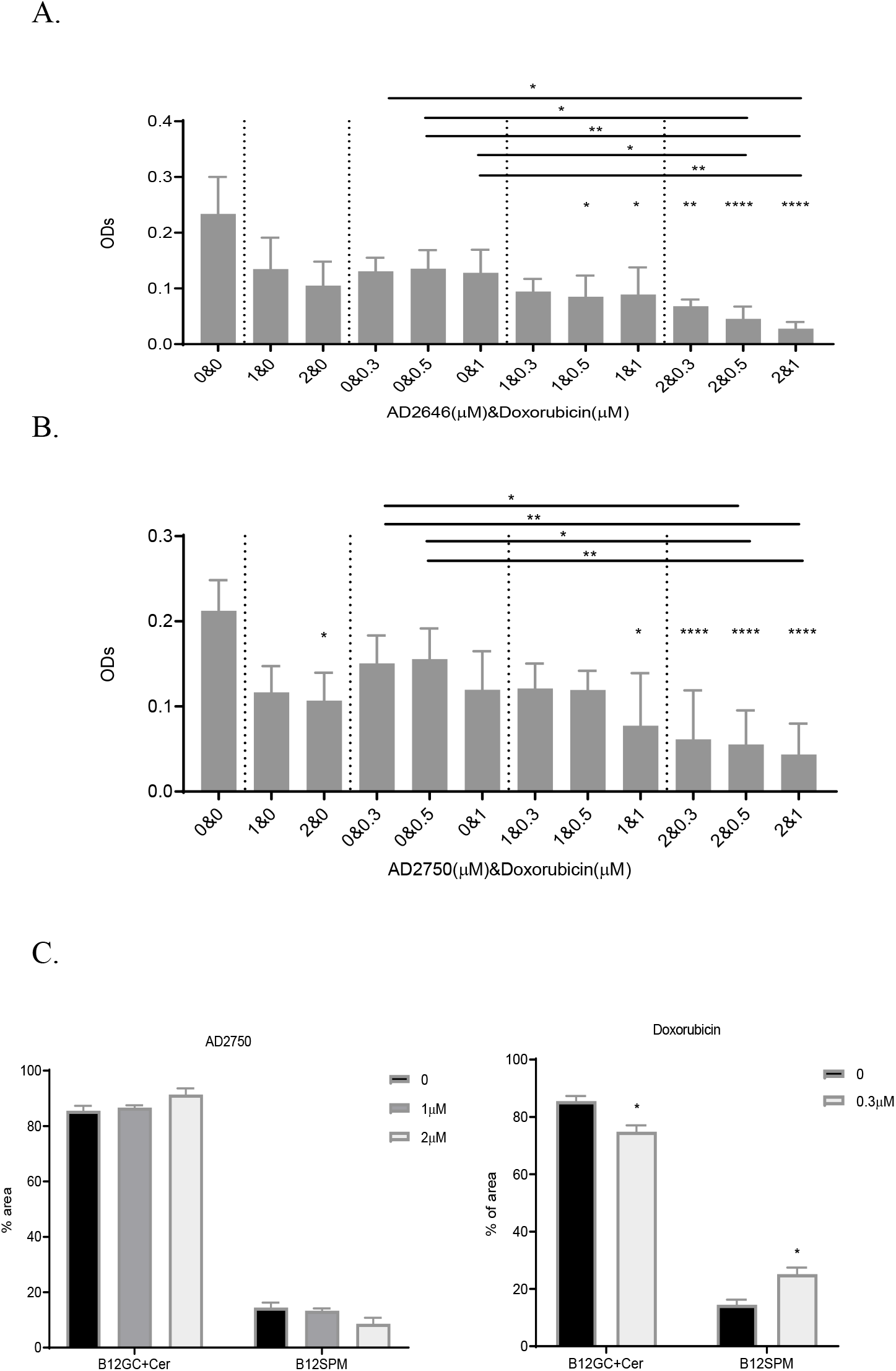
Doxorubicin and synthetic analogues synergistically increase cell death on human pancreatic cancer cells. CRL1687 cells were treated by the combination of (A) Doxorubicin (0.3μM, 0.5μM, 1μM) and AD2646 (1μM, 2μM) or (B) Doxorubicin (0.3μM, 0.5μM, 1μM) and AD2750 (1μM, 2μM) for 72 h. Cell viability was checked by MTT tests. ODs were shown. Kruskal-Wallis test was used, significance was compared to 0&0 unless being indicated. (C and D) Effects on the conversion of B12Cer+B12GC to B12SPM in CRL1687 extracts treated by (C) AD2750 (1μM, 2μM), and (D) Doxorubicin (0.3μM) are shown. CRL1687 cells were chase-labelled by BC12Cer for 30 minutes before adding analogues to cells culture for 3 days. Statistical significance determined using the Holm-Sidak method, with alpha = 0.05. Each experiment was repeated at least 3 times (Mean± SEM). Significance was compared to 0 unless being indicated. (*P<0.05, **P<0.01, ***P<0.001, ****P<0.0001.)

Taxol and AD2750 also synergistically reduced the cell viability of HT29 (Fig. 6A). When the cells were treated with Taxol (5μM) & AD2750 (2μM), or Taxol (5μM) & AD2750 (4μM), the cell viability decreased to 33% or 16.7%, respectively. When Taxol (5μM) or AD2646 (2μM, 4μM) were added to HT29, the cell viabilities were 74%, 67% and 37%, respectively. When the cells were treated with Taxol (5μM) & AD2646 (2μM), or Taxol (5μM) & AD2646 (4μM), it decreased to 37% or 14.8% (Fig. 6B). Adding Taxol and sphingolipid analogues to cells significantly reduced cell viability as compared to adding analogues individually.

**Figure.6.**
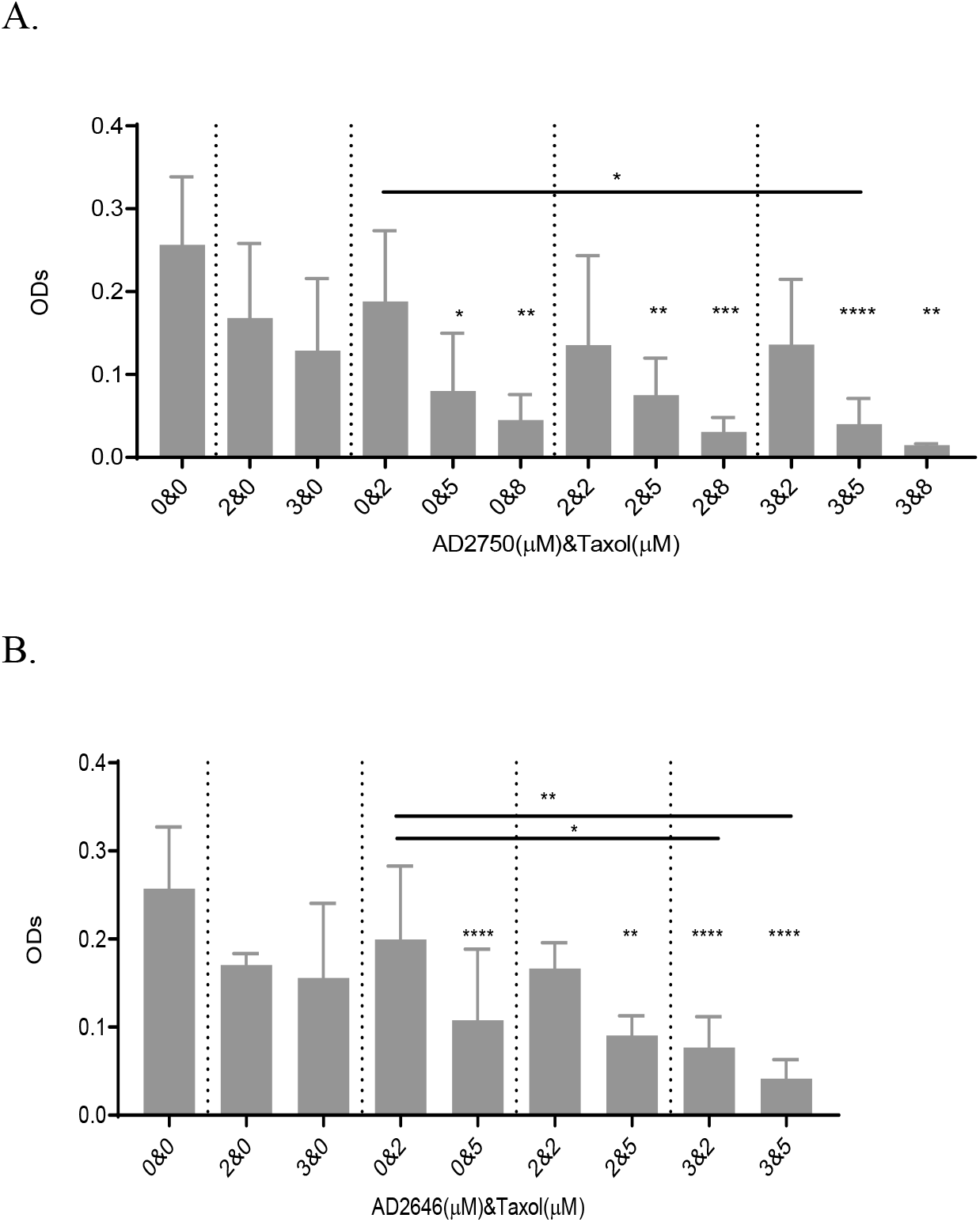
Taxol and synthetic analogues synergistically increase cell death on human colon cancer cells. HT29 cells were treated by (A) AD2750 (2μM, 3μM) and Taxol (2μM, 5μM, 8μM) or (B) AD2646 (2μM, 3μM) and Taxol (2μM, 5μM) for 72 h. Cell viability was checked by MTT tests. ODs were shown. Each experiment was repeated at least 3 times (Mean± SEM). Kruskal-Wallis test was used, significance was compared to 0&0 unless being indicated. (*P<0.05, **P<0.01, ***P<0.001, ****P<0.0001.)

### Taxol and AD2750 affect the metabolism of ceramide in HT29 cells

To understand whether Taxol enhances the effect of synthetic analogues on cell toxicity also through ceramide metabolism pathway, we labeled HT29 cells with B12Cer and then incubated cells with Taxol for 72h. Following the treatment with 0.04uM Taxol, the content of B12Cer increased from 40% to 90% and full conversion was observed at the concentration 0.1μM of Taxol, while no products of B12SPM and B12GC existed in the cell extractions. The content of B12GC was constantly around 6~8% under the treatment of different concentrations of Taxol (Fig 7A upper figure). Then, we tested the ceramide metabolism in cells with lower dose of Taxol (Fig 7A lower figure). It confirmed that Taxol could induce significant changes on the content of B12Cer and B12SPM at very low concentrations on HT29 cells. Even at 0.001μM, the amount of B12Cer and B12GC significantly increased compared to untreated cells, while B12SPM content dropped sharply (Fig 7A lower figure).

**Figure.7.**
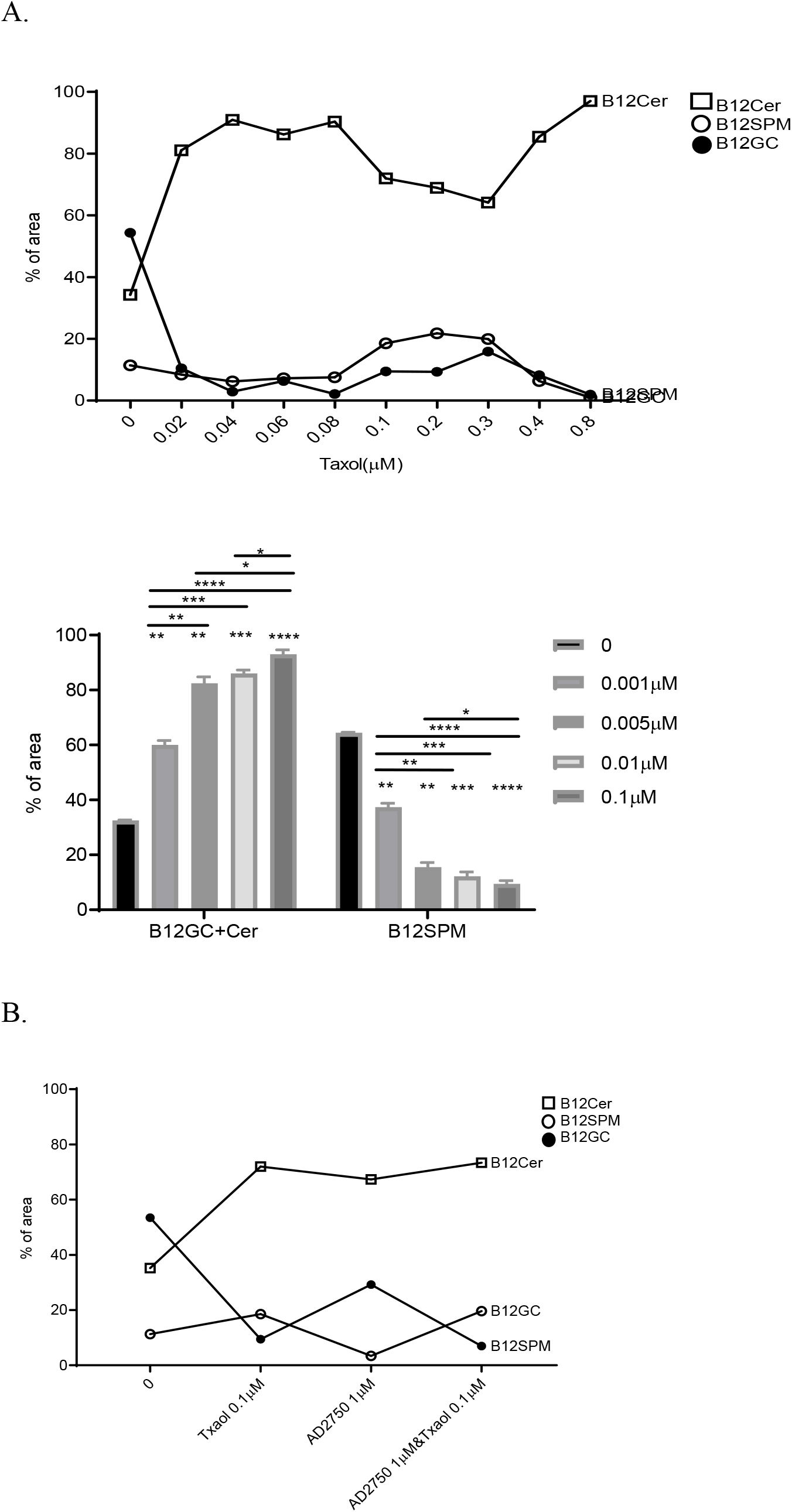
Taxol inhibits ceramide conversion in HT29. B12Cer labelled HT29 cells were incubated with (A) different concentrations of Taxol, or (B) Taxol (0.1μM), AD2750 (1μM) and Taxol (0.1μM) &AD2750 (1μM) for 72h, respectively. HT29 cell extraction was analyzed by HPLC. The square stands for B12Cer, the open circle stand for B12GC and the closed circle stands for B12SPM. Statistical significance determined using the Holm-Sidak method, with alpha = 0.05. Each experiment was repeated at least 3 times (Mean± SEM). Figure A (upper panel) and figure B experiments were done once. Significance was compared to 0 (untreated bar) unless being indicated. (*P<0.05, **P<0.01, ***P<0.001, ****P<0.0001.)

Since both compounds increased the content of ceramide in HT29 cells to induce cell death, we applied both compounds to HT29 cells and analyzed the cell extractions by HPLC. There was no obvious difference in the content of B12Cer and B12SPM on the cell extracts treated by the combination of Taxol (0.1μM) & AD2750 (1μM) in comparison with Taxol (0.1μM) or AD2750 (1μM) alone (Fig. 7B).

## Discussion

In this study, we show that synthesized sphingolipid analogues induced cell death on HT29, MDA-MB-435 and CRL1687 cells. No major effects were observed on ARH77 cells. Previously, we and others have shown that fluorescent lipids demonstrate the behavior of the naturally occurring ceramides (Morand, Fibach et al. 1982, Pagano and Sleight 1985, Dagan, Agmon et al. 2000). We also have shown that apoptosis induced by AD2646 through caspase-3 activation in Jurkat cells (Dagan, Wang et al. 2003, Qin, Weiss et al. 2010, Shimon Gatt 2012), and AD2900 and AD2750 induced apoptosis on human lymphocytes and inhibited the proliferation of human lymphocytes in mixed lymphocyte reaction (MLR) tests possibly through calcium pathway (Granot, Milhas et al. 2006, Song, Dagan et al. 2017). AD2730 and AD2673 also can induce apoptosis (Dagan, Wang et al. 2003, Shimon Gatt 2012). Due to the structural similarities of sphingolipid analogues to ceramide, sphingosine or trimethyl sphingosine (Dagan, Wang et al. 2003), our analogues can compete with and modify the metabolism of natural sphingolipids (Dagan, Wang et al. 2003, Darroch, Dagan et al. 2005, Granot, Milhas et al. 2006). We show that our analogues reduced the conversion of ceramide to sphingomyelin and glycolipids and in turn led to an increase of cellular ceramide in the fluorescence-labeled experiments, which then resulted in cell death possibly by apoptosis.

We show that all analogues influenced on the conversion of B12Cer below IC50s. Only AD2673, a sphingosine analog, behaves differently. AD2673 reduced glycolipid synthesis more pronounced than sphingomyelin synthesis (data not shown), and showed a higher IC50, affecting the metabolism of B12Cer to B12SPM under 10μM. Other publications have described syntheses and effects of a variety of analogues on metabolic pathways, especially inhibition of ceramidase, leading to an elevation of ceramide and resulting in apoptosis and cell death (Senchenkov, Litvak et al. 2001, Reynolds, Maurer et al. 2004, Ogretmen 2006, Hannun and Obeid 2008, Mao and Obeid 2008, Quinn, Wang et al. 2008). Thus our analogues could potentially affect two metabolic pathways, inhibiting the utilization of ceramide for biosynthesis of higher sphingolipids as well as the enzyme that hydrolyzes ceramide such as ceramidase (Mao and Obeid 2008, Shimon Gatt 2012). Each of these inhibitory effects elevated cell-ceramide, and a combination of the two elevated cell ceramide synergistically (Lucci, Han et al. 1999, Qiu, Zhou et al. 2006). Sphingolipid analogues inhibited ceramide metabolism, and then induced apoptosis might be valuable for potential applications as a therapeutic modality in cancer cells.

Taxol induces programmed cell death in leukemia and breast cancer cells. The co-administration of Taxol with exogenous ceramide substantially inhibits cell proliferation and elicits apoptosis in a synergistic fashion in Jurkat cells. Moreover, the effects of Taxol are linked to the *de novo* synthesis of ceramide in MDA-MB-468 and MCF-7 breast cancer cells and that Taxol-dependent cytotoxicity was abrogated by blocking ceramide production with L-cycloserine, an inhibitor of ceramide synthesis (Qiu, Zhou et al. 2006). In this study, Taxol significantly inhibited B12Cer conversion at very low concentrations on HT29 cells. When cells were challenged with sphingolipid analogues and very low dose of Taxol, ceramide can rise to toxic level, and then induced cell death. This could be due to the prominent inhibition effects of Taxol on ceramide conversion even at very low concentrations. Lower doses of Taxol or other possible pathway might need to be studied in the future. Low dose of Taxol and analogues may benefit cancer therapy.

Moreover, we have shown that the ceramide analogues AD2646 and AD2730 inhibited a purified acid ceramidase (Dagan, Wang et al. 2003). Together with the inhibition effects of our analogues on ceramide conversion, synthetic analogues are expected to increase the cellular levels of ceramide even more. Studies show that AD2750 can inhibit CRL1687 pancreatic cancer cell growth in CD1 nude mice (data not shown). In this study, we have proven that synergistic effects exist in a combination of Doxorubicin and AD2750 in human pancreatic cancer cells *in vitro*. In coordination, we would like to check this combination on the drug-resistance model. The synergistic effects of the combination of sphingolipid and commercial drugs might be of great importance in the treatment of some malignancies. Their anticancer activity should be tested by their application to *in vivo* tumor models in the future.

## Author contribution

JS and AD conceptualized and designed the study. JS was responsible for data collection. JS performed all experiments. JS wrote the manuscript with assistance from AD. AD obtained funding and was responsible for the entire project.

## Consent to Publish (Ethics)

All authors including Dr Jing Song and Dr. Arie Dagan, give our consent for the publication of identifiable details, which can include photograph(s) and/or videos and/or case history and/or details within the text (“Material”) to be published in the Molecular Biology Reports.

## Acknowledgements

We wish to thank to late Dr. Shimon Gatt for the support of this study.

## Figure legends

**Figure 1**. Structure of synthetic sphingolipid analogues.

**Figure 2**. B12SPM accumulated more in HT29. Different GC/SPM contents in human cancer cell extracts including HT29, MDA-MB-435, ARH77 and CRL1687. % of total area were shown. % of area stands for the percentage of area of each curve in total area of all curves in HPLC. The total of % of area is 1. Statistical significance determined using the Holm-Sidak method, with alpha = 0.05. Each experiment was repeated at least 3 times (Mean± SEM). (*P<0.05, **P<0.01, ****P<0.0001.)

**Figure.3**. Synthesized analogues inhibited the ceramide conversion on HT29 cells *in situ*. A). Retention times of B12GC, B12Cer, B12SPM from HT29 cell extracts are at 5.6, 5.9, 6.5 and 8.2minute analyzed by HPLC, respectively. HT29 cells were chase-labelled by BC12Cer and incubated for 24, 48, 72 or 96 h. B). Effects on the conversion of B12Cer to B12 GC and B12SPM in HT29 extracts treated by AD2646 (2μM, 4μM), AD2730 (0.8μM, 1.5μM, 2μM, 2.5μM), AD2750 (1μM, 2μM, 3μM), AD29000 (1μM, 2μM, 3μM) are shown. HT29 cells were labelled before adding analogues to cells culture for 3 days, % of total area was shown. Statistical significance determined using the Holm-Sidak method, with alpha = 0.05. Each experiment was repeated at least 3 times (Mean± SEM). Significance was compared to 0 unless being indicated. (*P<0.05, **P<0.01, ***P<0.001, ****P<0.0001)

**Figure.4**. Two synthetic analogues synergistically increase cell death on human cancer cells. A). MDA-MB-435 cells were treated by the combination of AD2900 (2.5μM, 3μM, 3.5μM) and AD2730 (1.5μM, 2μM, 2.5μM) for 24 h. B). HT29 cells were treated by the combination of AD2730 (3.5μM, 4μM, 4.5μM) and AD2750 (3.5μM, 4μM, 4.5μM) for 48 h. Cell viability was checked by MTT tests. ODs were shown. Each experiment was repeated at least 3 times (Mean± SEM). Kruskal-Wallis test was used, significance was compared to 0&0 unless being indicated. (*P<0.05, **P<0.01.)

**Figure.5**. Doxorubicin and synthetic analogues synergistically increase cell death on human pancreatic cancer cells. CRL1687 cells were treated by the combination of (A) Doxorubicin (0.3μM, 0.5μM, 1μM) and AD2646 (1μM, 2μM) or (B) Doxorubicin (0.3μM, 0.5μM, 1μM) and AD2750 (1μM, 2μM) for 72 h. Cell viability was checked by MTT tests. ODs were shown. Kruskal-Wallis test was used, significance was compared to 0&0 unless being indicated. (C and D) Effects on the conversion of B12Cer+B12GC to B12SPM in CRL1687 extracts treated by (C) AD2750 (1μM, 2μM), and (D) Doxorubicin (0.3μM) are shown. CRL1687 cells were chase-labelled by BC12Cer for 30 minutes before adding analogues to cells culture for 3 days. Statistical significance determined using the Holm-Sidak method, with alpha = 0.05. Each experiment was repeated at least 3 times (Mean± SEM). Significance was compared to 0 unless being indicated. (*P<0.05, **P<0.01, ***P<0.001, ****P<0.0001.)

**Figure.6**. Taxol and synthetic analogues synergistically increase cell death on human colon cancer cells. HT29 cells were treated by (A) AD2750 (2μM, 3μM) and Taxol (2μM, 5μM, 8μM) or (B) AD2646 (2μM, 3μM) and Taxol (2μM, 5μM) for 72 h. Cell viability was checked by MTT tests. ODs were shown. Each experiment was repeated at least 3 times (Mean± SEM). Kruskal-Wallis test was used, significance was compared to 0&0 unless being indicated. (*P<0.05, **P<0.01, ***P<0.001, ****P<0.0001.)

**Figure.7**. Taxol inhibits ceramide conversion in HT29. B12Cer labelled HT29 cells were incubated with (A) different concentrations of Taxol, or (B) Taxol (0.1μM), AD2750 (1μM) and Taxol (0.1μM) &AD2750 (1μM) for 72 h, respectively. HT29 cell extraction was analyzed by HPLC. The square stands for B12Cer, the open circle stand for B12GC and the closed circle stands for B12SPM. Statistical significance determined using the Holm-Sidak method, with alpha = 0.05. Each experiment was repeated at least 3 times (Mean± SEM). Figure A (upper panel) and figure B experiments were done once. Significance was compared to 0 (untreated bar) unless being indicated. (*P<0.05, **P<0.01, ***P<0.001, ****P<0.0001.)

**Supplementary figure 1**. IC50 (μM) of synthetic sphingolipid analogues AD2750 on human colon cancer cells (HT29) in MTT test. Experiments were done for 72 h. Each experiment was repeated at least 3 times.

**Supplementary figure 1.**
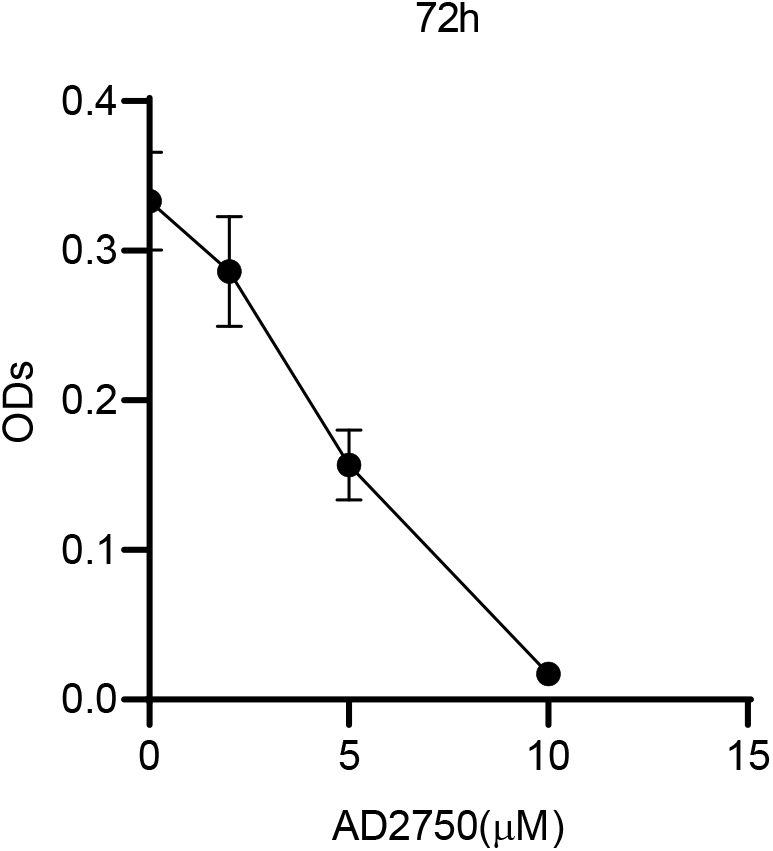
Curve of synthetic sphingolipid analogues AD2750 on human colon cancer cells (HT29) in a MTT test. Experiments were done for 72 h. Mean ± SEM was shown.

## References

Carvalho, C., R. X. Santos, S. Cardoso, S. Correia, P. J. Oliveira, M. S. Santos and P. I. Moreira (2009). Doxorubicin: the good, the bad and the ugly effect. Curr Med Chem. 16: 3267–3285;10.2174/092986709788803312.

Dagan, A., V. Agmon, S. Gatt and T. Dinur (2000). “Synthesis of fluorescent substrates and their application to study of sphingolipid metabolism in vitro and in intact cells.” Methods Enzymol 312: 293–30410.1016/s0076-6879(00)12916-7.

Dagan, A., C. Wang, E. Fibach and S. Gatt (2003). Synthetic, non-natural sphingolipid analogs inhibit the biosynthesis of cellular sphingolipids, elevate ceramide and induce apoptotic cell death. Biochimica et Biophysica Acta (BBA) - Molecular and Cell Biology of Lipids. 1633: 161–169;10.1016/s1388-1981(03)00122-7.

Darroch, P. I., A. Dagan, T. Granot, X. He, S. Gatt and E. H. Schuchman (2005). A lipid analogue that inhibits sphingomyelin hydrolysis and synthesis, increases ceramide, and leads to cell death. J. Lipid Res. 46: 2315–2324;10.1194/jlr.M500136-JLR200.

Fyrst, H. and J. D. Saba (2010). An update on sphingosine-1-phosphate and other sphingolipid mediators. Nat Chem Biol, Nature Publishing Group, a division of Macmillan Publishers Limited. All Rights Reserved. 6: 489–497;10.1038/nchembio.392.

Granot, T., D. Milhas, S. Carpentier, A. Dagan, B. Segui, S. Gatt and T. Levade (2006). Caspase-dependent and -independent cell death of Jurkat human leukemia cells induced by novel synthetic ceramide analogs. Leukemia. 20: 392–399;10.1038/sj.leu.2404084.

Guilbault, C., J. B. De Sanctis, G. Wojewodka, Z. Saeed, C. Lachance, T. A. A. Skinner, R. M. Vilela, S. Kubow, L. C. Lands, M. Hajduch, E. Matouk and D. Radzioch (2008). Fenretinide Corrects Newly Found Ceramide Deficiency in Cystic Fibrosis. Am. J. Respir. Cell Mol. Biol. 38: 47–56;10.1165/rcmb.2007-0036OC.

Haimovitz-Friedman, A., C. C. Kan, D. Ehleiter, R. S. Persaud, M. McLoughlin, Z. Fuks and R. N. Kolesnick (1994). Ionizing radiation acts on cellular membranes to generate ceramide and initiate apoptosis. The Journal of Experimental Medicine. 180: 525–535;10.1084/jem.180.2.525.

Hannun, Y. A. and L. M. Obeid (2008). Principles of bioactive lipid signalling: lessons from sphingolipids. Nat Rev Mol Cell Biol, Nature Publishing Group. 9: 139–150;10.1038/nrm2329

Labaied, M., A. Dagan, M. Dellinger, M. Gèze, S. Egée, S. L. Thomas, C. Wang, S. Gatt and P. Grellier (2004). Anti-Plasmodium activity of ceramide analogs. Malaria Journal. 3: 49;10.1186/1475-2875-3-49.

Lucci, A., T. Y. Han, Y. Y. Liu, A. E. Giuliano and M. C. Cabot (1999). Multidrug resistance modulators and doxorubicin synergize to elevate ceramide levels and elicit apoptosis in drug-resistant cancer cells. Cancer, John Wiley & Sons, Inc. 86: 300–311

Mao, C. and L. M. Obeid (2008). Ceramidases: regulators of cellular responses mediated by ceramide, sphingosine, and sphingosine-1-phosphate. Biochimica et Biophysica Acta (BBA) - Molecular and Cell Biology of Lipids. 1781: 424–434;10.1016/j.bbalip.2008.06.002.

Morad, S. A. F., T. S. Davis, M. Kester, T. P. Loughran and M. C. Cabot (2015). Dynamics of ceramide generation and metabolism in response to fenretinide – Diversity within and among leukemia. Leukemia Research. 39: 1071–1078;10.1016/j.leukres.2015.06.009.

Morand, O., E. Fibach, A. Dagan and S. Gatt (1982). “Transport of fluorescent derivatives of fatty acids into cultured human leukemic myeloid cells and their subsequent metabolic utilization.” Biochim Biophys Acta 711(3): 539–55010.1016/0005-2760(82)90070-4.

Ogretmen, B. (2006). Sphingolipids in cancer: Regulation of pathogenesis and therapy. FEBS Letters. 580: 5467–5476

Ogretmen, B. (2018). Sphingolipid metabolism in cancer signalling and therapy. Nature Reviews Cancer. 18: 33–50;10.1038/nrc.2017.96.

Pagano, R. E. and R. G. Sleight (1985). “Defining lipid transport pathways in animal cells.” Science 229(4718): 1051–105710.1126/science.4035344.

Qin, J. D., L. Weiss, S. Slavin, S. Gatt and A. Dagan (2010). “Synthetic, non-natural analogs of ceramide elevate cellular ceramide, inducing apoptotic death to prostate cancer cells and eradicating tumors in mice.” Cancer Invest 28(5): 535–54310.3109/07357900903478915.

Qiu, L., C. Zhou, Y. Sun, W. Di, E. Scheffler, S. Healey, H. Wanebo, N. Kouttab, W. Chu and Y. Wan (2006). Paclitaxel and ceramide synergistically induce cell death with transient activation of EGFR and ERK pathway in pancreatic cancer cells. Oncol Rep. 16: 907–913;10.3892/or.16.4.907.

Quinn, P. J., X. Wang, S. A. Saddoughi, P. Song and B. Ogretmen (2008). Roles of Bioactive Sphingolipids in Cancer Biology and Therapeutics. Lipids in Health and Disease, Springer Netherlands. 49: 413–440.

Radin, N. S. (2001). Killing cancer cells by poly-drug elevation of ceramide levels. European Journal of Biochemistry, Blackwell Science Ltd. 268: 193–204;10.1046/j.1432-1033.2001.01845.x.

Reynolds, C. P., B. J. Maurer and R. N. Kolesnick (2004). Ceramide synthesis and metabolism as a target for cancer therapy. Cancer Letters. 206: 169–180;10.1016/j.canlet.2003.08.034.

Senchenkov, A., D. A. Litvak and M. C. Cabot (2001). Targeting Ceramide Metabolismג€”a Strategy for Overcoming Drug Resistance. Journal of the National Cancer Institute. 93: 347–357;10.1093/jnci/93.5.347.

Shimon Gatt, A. D. (2012). Cancer and sphingolipid storage disease therapy using novel synthetic analogs of sphingolipids. Chemistry and Phsicys of Lipids. 165: 462–474;10.1016/j.chemphyslip.2012.02.006.

Song, J., A. Dagan, Z. Yakhtin, S. Gatt, S. Riley, H. Rosen, R. Or and O. Almogi-Hazan (2017). The novel sphingosine-1-phosphate receptors antagonist AD2900 affects lymphocyte activation and inhibits T-cell entry into the lymph nodes. Oncotarget, Impact Journals LLC. 8: 53563–53580;10.18632/oncotarget.18626.

Swanton, C., M. Marani, O. Pardo, P. H. Warne, G. Kelly, E. Sahai, F. d. r. Elustondo, J. Chang, J. Temple, A. A. Ahmed, J. D. Brenton, J. Downward and B. Nicke (2007). Regulators of Mitotic Arrest and Ceramide Metabolism Are Determinants of Sensitivity to Paclitaxel and Other Chemotherapeutic Drugs. Cancer Cell. 11: 498–512;10.1016/j.ccr.2007.04.011.

Testi, R. (1996). Sphingomyelin breakdown and cell fate. Trends in Biochemical Sciences. 21: 468–471;10.1016/s0968-0004(96)10056-6.

Vethakanraj, H. S., B. P. Sesurajan, V. P. Padmanaban, M. Jayaprakasam, S. Murali and A. K. Sekar (2018). Anticancer effect of acid ceramidase inhibitor ceranib-2 in human breast cancer cell lines MCF-7, MDA MB-231 by the activation of SAPK/JNK, p38 MAPK apoptotic pathways, inhibition of the Akt pathway, downregulation of ERα. Anti-cancer drugs. 29: 50–60;10.1097/cad.0000000000000566.

Wang, H., B. J. Maurer, Y.-Y. Liu, E. Wang, J. C. Allegood, S. Kelly, H. Symolon, Y. Liu, A. H. Merrill, V. r. Gouaz©-Andersson, J. Y. Yu, A. E. Giuliano and M. C. Cabot (2008). N-(4-Hydroxyphenyl)retinamide increases dihydroceramide and synergizes with dimethylsphingosine to enhance cancer cell killing. Molecular Cancer Therapeutics. 7: 2967–2976;10.1158/1535-7163.mct-08-0549

